# Oocyte and cumulus cell cooperativity and metabolic plasticity under the direction of oocyte paracrine factors

**DOI:** 10.1101/2022.09.05.506599

**Authors:** Dulama Richani, Anne Poljak, Baily Wang, Saabah B. Mahbub, Joanna Biazik, Jared M. Campbell, Abbas Habibalahi, William A. Stocker, Maria B. Marinova, Brett Nixon, Sonia Bustamante, David Skerrett-Byrne, Craig A. Harrison, Ewa Goldys, Robert B. Gilchrist

## Abstract

Mammalian oocytes develop and mature in a mutually dependent relationship with surrounding cumulus cells. The oocyte actively regulates cumulus cell differentiation and function by secreting soluble paracrine oocyte-secreted factors (OSFs). We characterized the molecular mechanisms by which two model OSFs, cumulin and BMP15, regulate oocyte maturation and cumulus-oocyte cooperativity. Exposure to these OSFs during maturation altered the proteomic and multispectral autofluorescence profiles of both the oocyte and cumulus cells. In oocytes, cumulin significantly upregulated proteins involved in nuclear function. In cumulus cells, both OSFs elicited marked upregulation of a variety of metabolic processes (mostly anabolic), including lipid, nucleotide, and carbohydrate metabolism, while mitochondrial metabolic processes were downregulated. The mitochondrial changes were validated by functional assays confirming altered mitochondrial morphology, respiration, and content, whilst maintaining ATP homeostasis. Collectively, these data demonstrate that OSFs remodel cumulus cell metabolism during oocyte maturation in preparation for ensuing fertilization and embryonic development.

**HIGHLIGHTS:**

- During oocyte maturation, oocyte-secreted factors promote cell cooperativity between the oocyte and cumulus cells by altering the molecular composition of both cell types.
- Oocyte-secreted factors downregulate protein catabolic processes, and upregulate DNA binding, translation, and ribosome assembly in oocytes.
- Oocyte-secreted factors alter mitochondrial number, morphology, and function in cumulus cells.
- Oocyte-secreted factors further enhance metabolic plasticity in cumulus cells by upregulating anabolic pathways for macromolecules and small molecule organics.
- The oocyte, via oocyte-secreted factors, instructs cumulus cells to increase metabolic workload on its behalf, thereby subduing oocyte metabolism.

## INTRODUCTION

While it is a fundamental tenet of biology that the basic unit of life is a cell, however complex, multicellular life would not be possible without symbiotic relationships amongst cells of quite different phenotype. Symbiotic relationships are prevalent throughout nature and are conventionally defined as relationships between different organisms (e.g., gut microbiota and the host organism). However, “auto-symbiosis” or cellular cooperativity, is also typical of most, if not all, organ systems within multicellular organisms [1-3].It is a likely mechanism/driver of evolution of eukaryotic cells [4] and is a common state of cellular systems organisation within all multicellular life. Here we focus on the mechanisms that regulate a cooperative inter-cellular relationship which precedes and underpins sex-based life in animals; germ-somatic cell cooperativity (auto-symbiosis).

Mammalian germ cells develop and mature in an intimate and mutually dependent relationship with adjacent somatic cells. The auto-symbiotic relationship between the female germ cell, the oocyte, and its somatic cells, is unique and essential as the oocyte is a highly unusual cell, being the largest cell in the body, long-lived, meiotically arrested, and metabolically inept. As such it is entirely dependent on its support somatic cells for survival and appropriate development. By the end of the oocyte growth phase, the oocyte is surrounded by cumulus cells, which nurture the oocyte through its final phases of development, including the meiotic maturation phase immediately prior to ovulation. Such support is essential to the subsequent developmental capacity of the oocyte, as oocytes matured in the absence of their cumulus vestment exhibit a lower capacity to support subsequent embryo development [5].Cumulus cells form an intimate physical association with the oocyte via highly specialized cytoplasmic transzonal projections, which penetrate through the oocyte’s zona pellucida, forming a structure called the cumulus–oocyte complex (COC) [6]. Gap junctions are at the termini of these projections and facilitate the transfer of small regulatory factors and metabolites to the oocyte, which are essential for oocyte development.

This oocyte-cumulus cell association has multiple advantages for maintaining genomic stability and integrity of the oocyte, which is central to reproductive success, by; ***(a)*** protecting oocytes from the stress of external impacts, and mediating environmental signals on behalf of the oocyte, and ***(b)*** providing essential nutrients and metabolites, minimizing the metabolic demands on the oocyte, thereby also minimizing secondary production of potentially DNA damaging free radicals. Hence, in many respects, the oocyte is reliant on cumulus cells for its normal function. Consequently, it was long thought that the mammalian oocyte is passive in its relationship with cumulus cells, but it is now clear that the communication axis is birather than mono-directional. Over the last two decades a new paradigm has emerged that the oocyte is in fact central to regulating the differentiation and critical functions of cumulus cells, which it achieves via the secretion of paracrine growth factors [5, 7-9]. For example, cumulus cells are not metabolically competent in the absence of an oocyte, exhibiting perturbations in metabolically driven processes, such as cumulus expansion [10-13]. Oocytes direct appropriate gene expression, potently stimulate DNA synthesis and cellular proliferation, prevent apoptosis, maintain cellular differentiation, and stimulate glycolysis and amino acid transport in cumulus cells (reviewed by Gilchrist, Lane [14]). Such regulation of cumulus cell function in turn affects the oocyte’s own development and its capacity to support subsequent embryo and fetal development [15-17].

Growth differentiation factor-9 (GDF9) and bone morphogenetic protein-15 (BMP15) are oocyte-secreted members of the TGF-β superfamily, identified as central regulators of cumulus cell differentiation [18-20]. Genetic, biochemical and protein functional studies have demonstrated potent synergistic interactions between GDF9 and BMP15 [21]. In particular, the GDF9:BMP15 heterodimer, cumulin, has potent activity relative to GDF9 and BMP15 homodimers [22-25]. In vitro exposure of COCs to cumulin promotes greater cell proliferation and cumulus expansion and improves subsequent embryo development in murine, porcine, and equine models [24, 26, 27]. Thus, cumulin has potential as an in vitro maturation (IVM) supplement to improve assisted reproductive technologies (ART), such as IVF, used to treat infertility.

Although there is no doubt that the oocyte actively regulates cumulus cell function using oocyte-secreted factors (OSFs) [14, 28], the somatic cellular processes regulated by OSFs, that in turn impact the developmental program of the germ cell, are still emerging. Here we investigate the molecular mechanisms by which the OSFs, cumulin and BMP15, regulate oocyte maturation and their effect on cumulus-oocyte cooperativity. Global analyses (proteomics and multispectral analysis) reveal proteomic and metabolic profiles which discriminate cumulin and BMP15 treated cells from controls and from each other, and reveal the distinct molecular pathways triggered within each cell type in this auto-symbiotic relationship. Targeted analyses indicate metabolic plasticity which redirects mitochondrial metabolism towards a massive increase of cytoplasmic anabolic pathways in cumulus cells, which subsume multiple “housekeeping” roles on behalf of oocytes. This inter-cellular cooperativity facilitates oocyte maturation while simultaneously protecting germ-line genomic integrity, in a manner which could not be achieved by a single cell alone.

## MATERIALS & METHODS

### Recombinant cumulin and BMP15 production

Recombinant human cumulin and human BMP15, both as purified pro-mature complexes, were used in this study. As both BMP15 and GDF9 normally exist as non-covalent dimers, covalent dimers were generated by substituting a serine residue with a cysteine (BMP15^S356C^ and GDF9^S418C^, respectively), enabling the formation of the inter-subunit disulphide bond which stabilizes most other TGF-β superfamily ligands, preventing dimer dissociation [23, 29]. Production of recombinant cumulin (batch 8a) and BMP15 (batch 7a) was carried out in-house, as described previously [22, 23]. Briefly, HEK293T cells were transfected with expression plasmids for human BMP15^S356C^ ± GDF9^S418C^ to produce either BMP15 or cumulin, respectively, before being placed into production medium (DMEM:F-12 medium containing 0.02% BSA and 0.005% heparin). Protein purification was carried out by eluting bound proteins from cobalt-based immobilised metal affinity chromatography resin with elution buffer (50 mM phosphate buffer, 300 mM NaCl, 200 mM imidazole, pH 7.4) followed by dialysis against phosphate buffer to remove imidazole [22, 23].

### Oocyte in vitro maturation (IVM)

Mice were maintained in accordance with the Australian Code of Practice for Care and Use of Animals for Scientific Purposes and all experimental protocols were approved by the University of New South Wales Sydney Animal Care & Ethics Committee (ethics 17/105A). Peri-pubertal 28-30-day old C57/Bl6 females were given an intraperitoneal injection of 5 IU of equine chorionic gonadotropin (eCG; Folligon, Intervet, Boxmeer, The Netherlands). At 46 h post-eCG, ovaries were harvested and placed in HEPES-buffered alpha minimum essential medium (αMEM; Gibco, Life Technologies, New York, USA) supplemented with 3 mg/mL BSA (MP Biomedicals, Auckland, NZ). COCs were released from preovulatory follicles using a 27-gauge needle into HEPES-buffered αMEM (Gibco) with 3 mg/mL of bovine serum albumin (BSA) and 100 µM of IBMX (Sigma-Aldrich, Merck, Darmstadt, Germany), and collected. COCs were washed 3 times and then cultured for 17 h in bicarbonate-buffered αMEM (Gibco) supplemented with 3 mg/mL BSA, mouse amphiregulin and epiregulin (50 ng/mL each; R&D Systems, Minneapolis, USA), +/-cumulin (20 ng/mL) or +/-BMP15 (20 ng/mL). Where oocytes and cumulus cells were analyzed separately, oocytes were denuded after 17 h of IVM by mechanical shearing using a P200 pipette. A diagrammatic illustration of experimental design, cell types analyzed and end-points assessed is provided in Figure 1.

**Figure 1:**
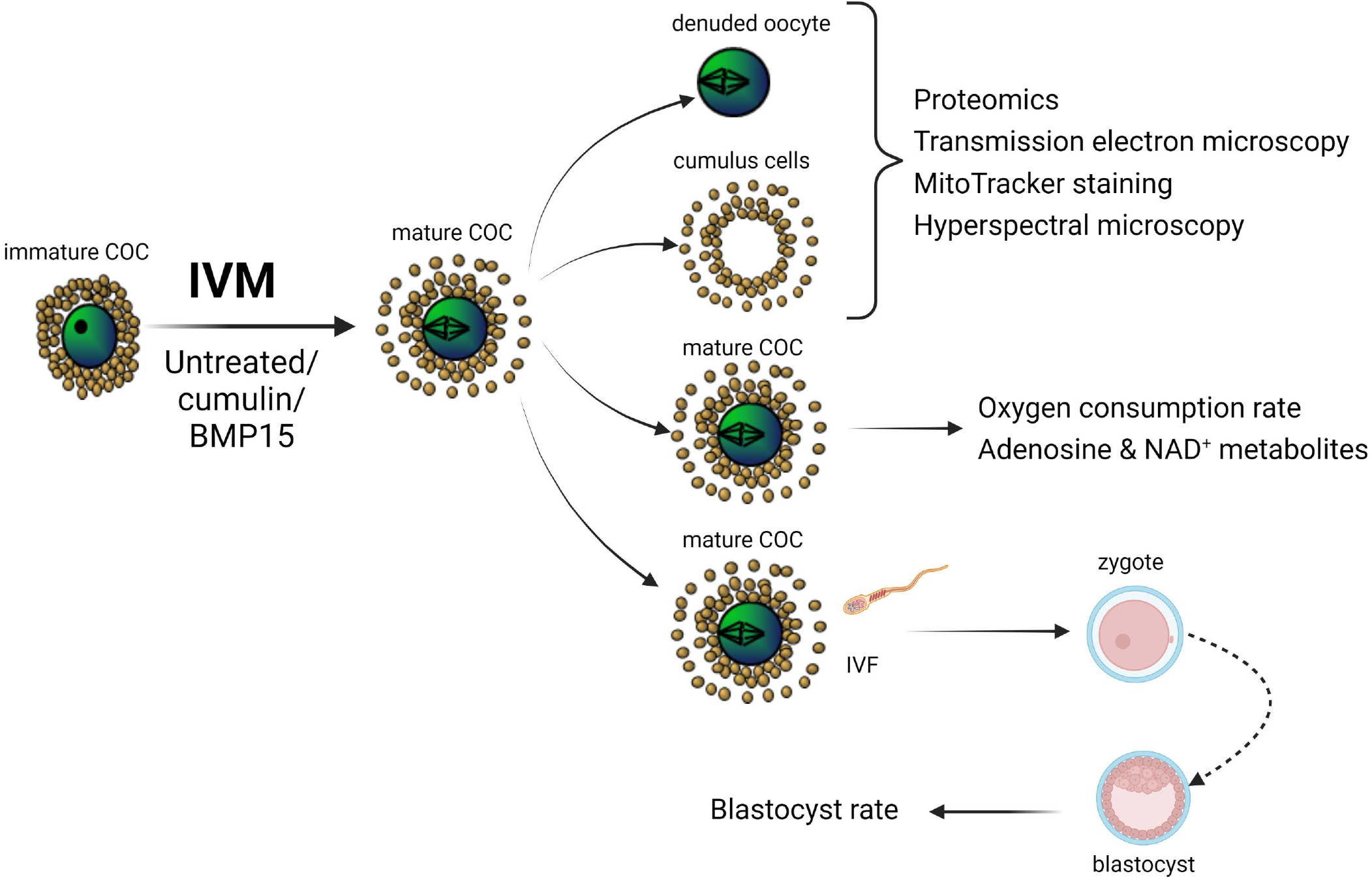
Schematic of study experimental design and endpoints. Immature mouse cumulus-oocyte complexes (COCs) underwent in vitro maturation (IVM) (17 h of culture ± cumulin or BMP15 in the culture medium). Following IVM, mature COCs were either analyzed whole, or oocytes and surrounding cumulus cells were separated and analyzed separately. IVF, in vitro fertilization.

### In vitro embryo production

Following IVM, COCs were fertilized and embryos cultured for 6 days as previously described by Stocker, Walton [24]. All media were purchased from IVF Vet Solutions (Adelaide, Australia). Following IVM as described above, COCs were washed (Research Wash, supplemented with 4 mg/mL BSA) and co-incubated with CBB6F1 capacitated sperm for 3.5 h at 37°C with 5% O_2_, 6% CO_2_ and 89% N_2_ in fertilization medium (Research Fert) with 4 mg/mL BSA. Presumptive zygotes were cultured in cleavage medium (Research Cleave) with 4 mg/mL BSA. Embryo development was assessed every 24 h over 6 days.

### Protein extraction and mass spectrometry for proteomic analysis

Four biological replicates were included for each of control and cumulin treated cells, and three biological replicates for BMP15 treated cells. Full mass spectrometry methodology and bioinformatics analysis is provided in the supplementary methods. Briefly, cells were lysed and protein extracted. Samples were analyzed using a QExactive mass spectrometer (Thermo Electron, Bremen, Germany). Peptide separation was carried out by nano-liquid chromatography (nano-LC) on a Dionex UltiMate 3000 HPLC system (ThermoScientific, Waltham, MA), equipped with an autosampler (Dionex, Amsterdam, Netherlands).

### Bioinformatics analysis

In this study, data were processed using search engine algorithms for protein identification and abundance quantification (Mascor+Scaffold and ProteomeDiscoverer v2.4). The results were merged based on consistency between direction of treatment/control ratio change. A further requirement was that at least one, if not both methods, had a p-value of <0.05. Basic parameter settings kept common across workflows were: taxonomy *Mus musculus*, database Uniprot, enzyme trypsin, maximal missed cleavages = 2, MS1 tolerance = ±10ppm, MS2 tolerance = ±0.05Da, fixed modification carbamidomethyl (C), variable modifications acetyl (N-terminal), oxidation (M), and phosphorylation (S,T,Y). Proteins were identified with a minimum of 2 peptides each. Both peptide and protein false discovery rate (FDR) using the decoy database data were <1%. Relative quantification was based on normalized spectral abundance factor (NSAF) ratios of treatment vs control samples, and a *t*-test was used to determine statistically significant expression differences between treatment and control groups [30, 31]. The differentially expressed proteins list was further analyzed using STRING software to explore clustering and enrichment of specific molecular functions, and biological pathways. Detailed methodology and rationale for this approach is provided in the supplementary methods.

### Transmission electron microscopy (TEM)

Following IVM, oocytes were fixed at 4°C overnight in a fixative comprising 2.5% w/v-1 glutaraldehyde in 0.2 M sodium cacodylate buffer. Fixed oocytes were rinsed with 0.1 M sodium cacodylate buffer and post fixed in 1% osmium tetroxide in 0.2 M sodium cacodylate buffer by using a BioWave Pro+ Microwave Tissue Processor (Ted Pella, USA). After rinsing with 0.1 M sodium cacodylate buffer, oocytes were dehydrated with a graded series of ethanol, infiltrated with resin (Procure, 812), and polymerized using an oven at 60°C for 48 h. Ultrathin sections (70 nm) were cut using a diamond knife (Diatome) and collected onto carbon-coated copper slot TEM grids. Grids were post-stained using uranyl acetate (2%) and lead citrate. Duplicates from each of the three treatments (n=16 cumulus cells, n=6 oocytes) were imaged using a JEOL 1400 TEM (Tokyo, Japan) operating at 100 kV.

### COC oxygen consumption

Following IVM, COC oxygen consumption rate was measured using a Seahorse XFe96 platform (Agilent, Santa Clara, CA, USA). Sensor-containing Seahorse XFe96 cartridges (Agilent) were hydrated with sterile ultrapure water overnight at 37°C as per manufacturer’s instructions. Following IVM, COCs were placed into 96-well plates (previously coated with 22.4 µg/mL Cell-Tak) containing pre-warmed XF DMEM supplemented with 1 mM pyruvate, 2 mM glutamine and 5 mM glucose, and allowed to equilibrate at 37°C in ambient air for 1 h. COCs were analyzed using the Seahorse XFe96 platform with a Seahorse XF Mito Stress test kit (to assess mitochondrial oxygen consumption) as per the manufacturer’s instructions. Final concentrations of mitochondrial inhibitors used were 1 µM oligomycin, 2.5 µM FCCP, and 2.5 µM rotenone + antimycin A (Agilent). Following the completion of each assay, COCs per well were visualized to ensure they remained adhered to the bottom of the well. Data were normalized to COC number, with each well containing 5-15 COCs.

### Multispectral microscopy

Multispectral imaging of oocyte and cumulus cell autofluorescence were conducted on an Olympus IX83 epifluorescent microscope with LED illumination at various excitation/emission pairs (listed in supplementary); where excitation values are ± 5 nm and emissions are ± 20nm. Unmixing of specific fluorophores was carried out by linear mixed modelling (LMM) in which extracted spectral characteristics were compared to the known characteristics of fluorophores. Discrimination between groups, whether cumulus cells or oocytes, was performed using a variety of quantitative cellular image features including mean channel intensity, channel intensity ratio [32], color distribution [33], textural features [34] and others defined elsewhere [32-34]. Candidate features (selected if p<0.005 by ANOVA) were projected onto an optimal two-dimensional space created by discriminative analysis [35, 36]. A linear classifier was applied based on a linear predictor function that utilized a set of weights obtained from assessment during a training process [37, 38]. Full methodology is given in the supplementary section.

### Metabolite analysis

Liquid chromatography with tandem mass spectrometry (LC-MS/MS) was used to quantify energy nucleotides in COCs across treatment conditions, based on a previously published approach [39, 40]. A detailed explanation of the method is provided in supplementary methods.

### MitoTracker staining

MitoTracker Orange, a fluorescent dye which stains mitochondria and whose accumulation is dependent on membrane potential, was used to estimate mitochondrial numbers across treatment conditions. Following IVM, COCs were incubated with MitoTracker Orange (ThermoFisher Scientific, diluted to 200 nM in HEPES-buffered αMEM) for 15 mins. COCs were washed in medium and placed in 4% paraformaldehyde (in 0.2 M Na_3_PO_4_ buffer) for 1 h at room temperature. COCs were then placed on glass-bottom dishes in 0.1 M Na_3_PO_4_ buffer overlaid with oil and stored at 4°C in the dark for up to 1 week before confocal imaging. The COCs were imaged using a Zeiss LSM 880 with LD C-Apochromat 40X/1.1 W Korr M27 objective. One section was taken though the middle of the oocyte in 8-bit image depth with 1024 × 1024 pixel image resolution and pixel size of 0.21 µm. The cells were irradiated with a 561 nm laser and emission was detected in the 566-684 nm range. Due to the brightness difference between the oocyte and cumulus cells, two different exposure settings were used. Each COC was imaged twice with exposures optimal for oocyte and cumulus cells. Those settings were kept constant across all repeats.

### Statistical analyses

Blastocyst rates (total and hatching) were calculated as a proportion of cleaved embryos. Blastocyst rates were represented in percentages and arcsine transformed prior to one-way ANOVA with Tukey’s post-hoc test for statistical analysis. MitoTracker staining and data from metabolite analysis by mass spectrometry were analysed by one-way ANOVA with Tukey’s (parametric data) or Kruskal-Wallis (non-parametric data) post-hoc tests. Effect of treatments on mitochondrial OCR was analysed by Mann-Whitney test. Data were analysed using GraphPad Prism 9.3.1. The relative abundance of metabolites analysed by multispectral imaging were analysed by two-sample t-tests using Matlab 2020a. Statistical significance was considered at p≤0.05.

## RESULTS

Assessment of oocyte and cumulus cell proteomic and multispectral profiles, cell morphology, metabolism, and subsequent embryo development, were assessed as outlined in Fig. 1.

### Cumulin improves oocyte developmental competence

Embryo development was examined to assess whether the developmental competence of COCs matured via IVM is enhanced by cumulin. COCs treated with increasing doses of cumulin were fertilized and their capacity to support embryo development to the blastocyst stage was assessed (Fig. 2A). Relative to untreated controls, oocytes matured in the presence of 20 ng/mL cumulin did not exhibit significantly altered day 5 blastocyst yield (Fig. 2B), however significantly higher blastocyst (Fig. 2C) and hatching blastocyst yields (Fig. 2D) was observed on day 6, by ∼30% in each case.

**Figure 2:**
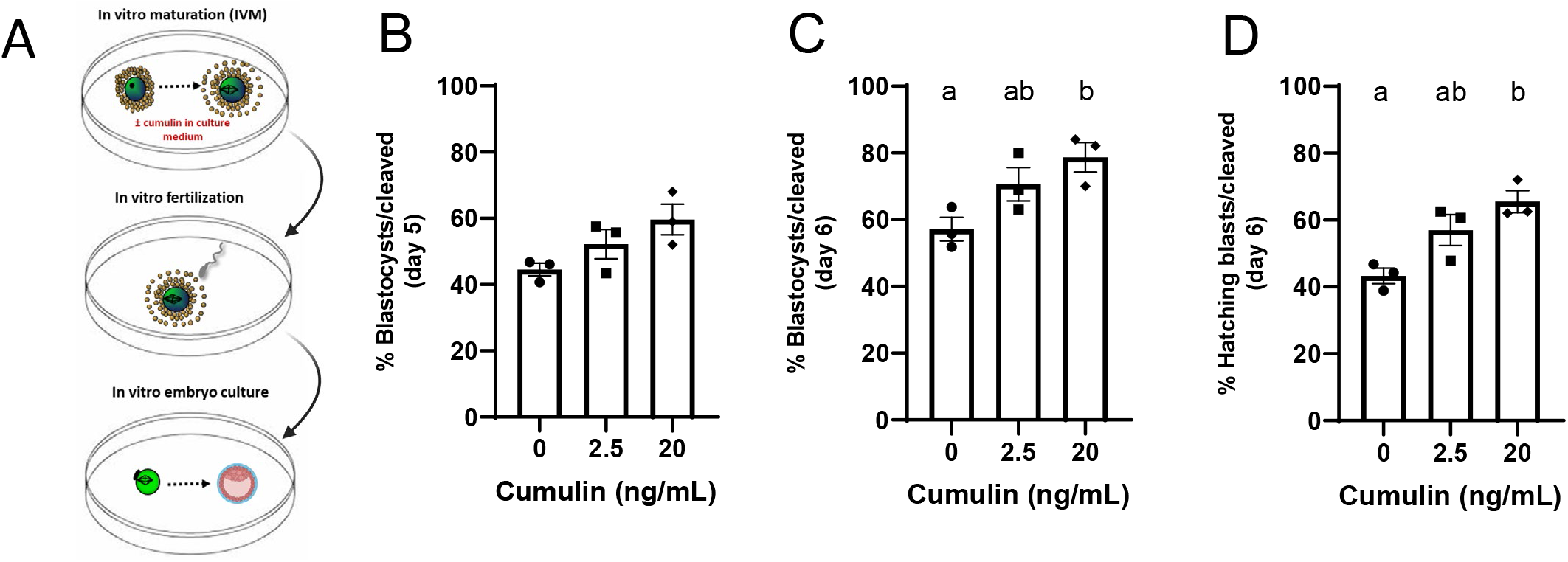
(A) Effect of cumulin on embryo development (as a marker of oocyte developmental competence) following 17 h of IVM. (B) Day 5 total blastocyst, (C) day 6 total blastocyst, (D) day 6 hatching blastocyst rates were assessed over 3 biological replicates. Bars represent the mean ± SEM, with 162-180 oocytes used per treatment group. Data were arcsine transformed and one-way ANOVA followed by Tukey’s post-hoc tests were performed. Bars with no common superscripts are significantly different (p<0.05).

### Global analyses using proteomics and hyperspectral analysis

Global proteomic and multispectral analyses of oocytes and cumulus cells post-IVM are shown in Figure 3 (full protein lists are shown in Supplementary Tables S1 and S2). Cell types and treatment conditions can be clearly distinguished based on these orthogonal global approaches. Overall, more proteins and more differentially expressed proteins were identified in cumulus cell samples compared to oocytes (Fig. 3A and 3B), due, at least in part, to the considerably larger amount of total protein available in the cumulus cell samples (15 µg for cumulus cells vs 2-3 µg for oocytes). Proteomic heat maps (Fig. 3C) and multispectral analysis plots (Fig. 3D) both show distinct profiles, with limited overlap between controls, BMP15 and cumulin treated COCs, in both cell types. A more detailed explanation of the proteomic results relating to spectral counting (Scaffold) and peak area integration (PD2.4) is provided in the Supplementary Results.

**Figure 3:**
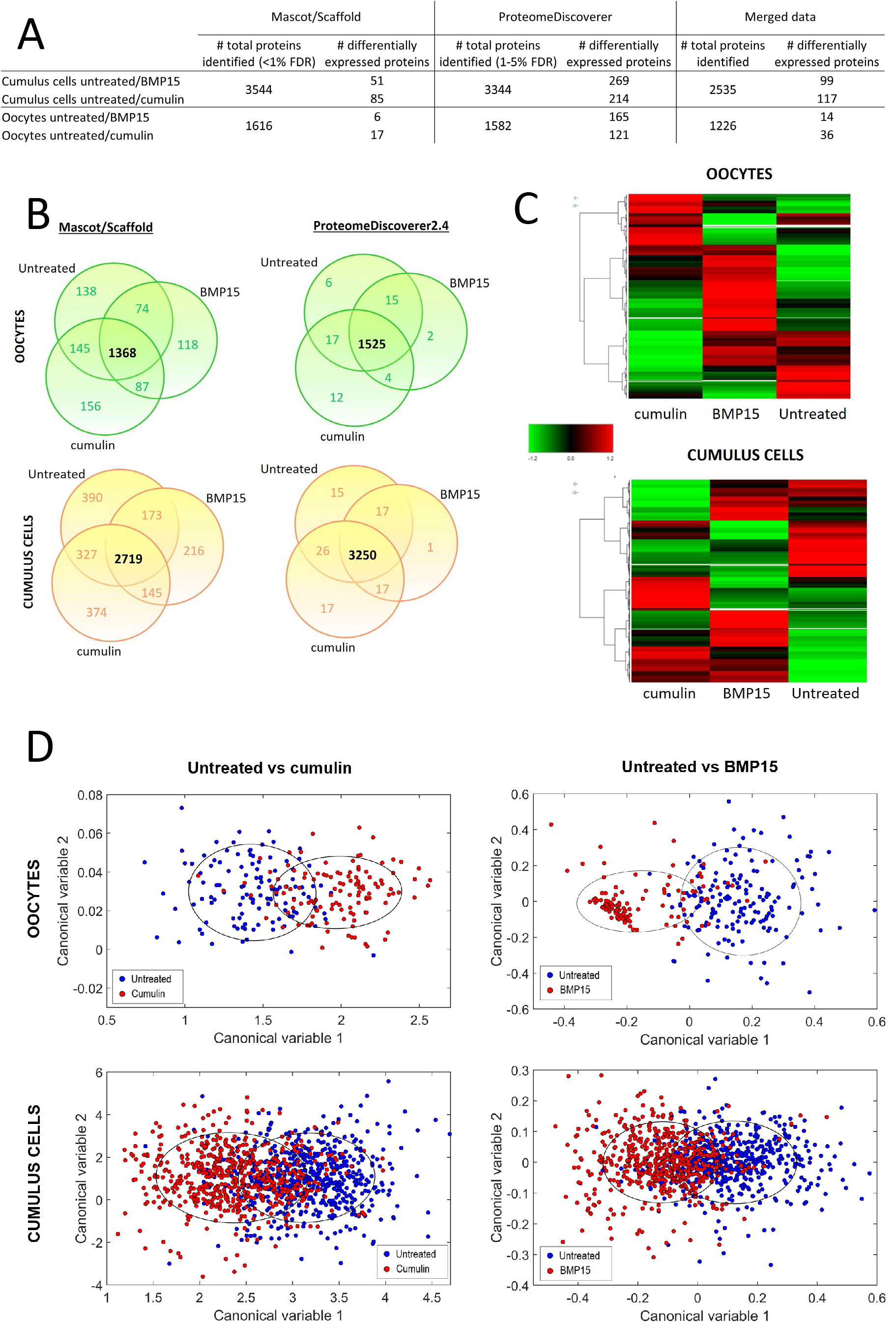
Global analyses of proteomic and multispectral intensity change in BMP15 and cumulin treated COCs. (A) Total number of proteins identified and number of differentially expressed proteins between treatment groups. The merged dataset represents proteins confidently identified in both workflows. The significantly different proteins under the merged column are those proteins whose ratios of expression change in the same direction in both workflows and are significantly different to controls in at least one workflow. (B) Treatment group overlaps of numbers of proteins identified in each of two search engines (Mascot/Scaffold and ProteomeDiscoverer2.4). (C) Heat maps of protein abundance distribution across treatment groups. (D) Linear discriminant analysis of 10-20 multispectral signature features that varied significantly (p<0.005) between cumulus cells and oocytes in response to cumulin or BMP15. N= 115-158 oocytes and n=532-600 cumulus cells analyzed per treatment group. Blue circles = untreated cells, red circles = treated cells.

### Effects of BMP15 and cumulin treatment of COCs on oocyte proteome expression

More than double the number of proteins were differentially expressed in oocytes following cumulin treatment of COCs, as compared with BMP15 treatment (Figs. 3A, 4A). In oocytes, fewer of the differentially expressed proteins were downregulated, with the majority being upregulated in response to both treatments (Fig. 4A). Network analysis of differentially expressed proteins showed relatively few significantly enriched networks in response to BMP15 (7 in total), with protein folding and nucleosome being the two main upregulated networks, and no significantly enriched networks apparent in the short list of downregulated proteins (Figs. 4A, 4C, 4E). By contrast, cumulin induced a diverse range of significantly enriched networks in oocytes (37 and 60 respectively, in the up- and down-regulated protein lists, Figs. 4A, 4C, 4E). Significant networks of upregulated proteins in oocytes included cell envelope and cytoskeletal modification/organization of organelles, oxidoreductase activity, DNA binding and ribosomal constituents (Fig. 4A, 4C), while significant networks of downregulated proteins included protein complex organization, ribonucleotide binding and intracellular organelles. Interestingly, even though there were a greater number of upregulated than downregulated proteins in oocytes following cumulin treatment, there were fewer networks amongst the upregulated proteins, and almost a third more significant networks amongst the shorter downregulated protein list.

**Figure 4:**
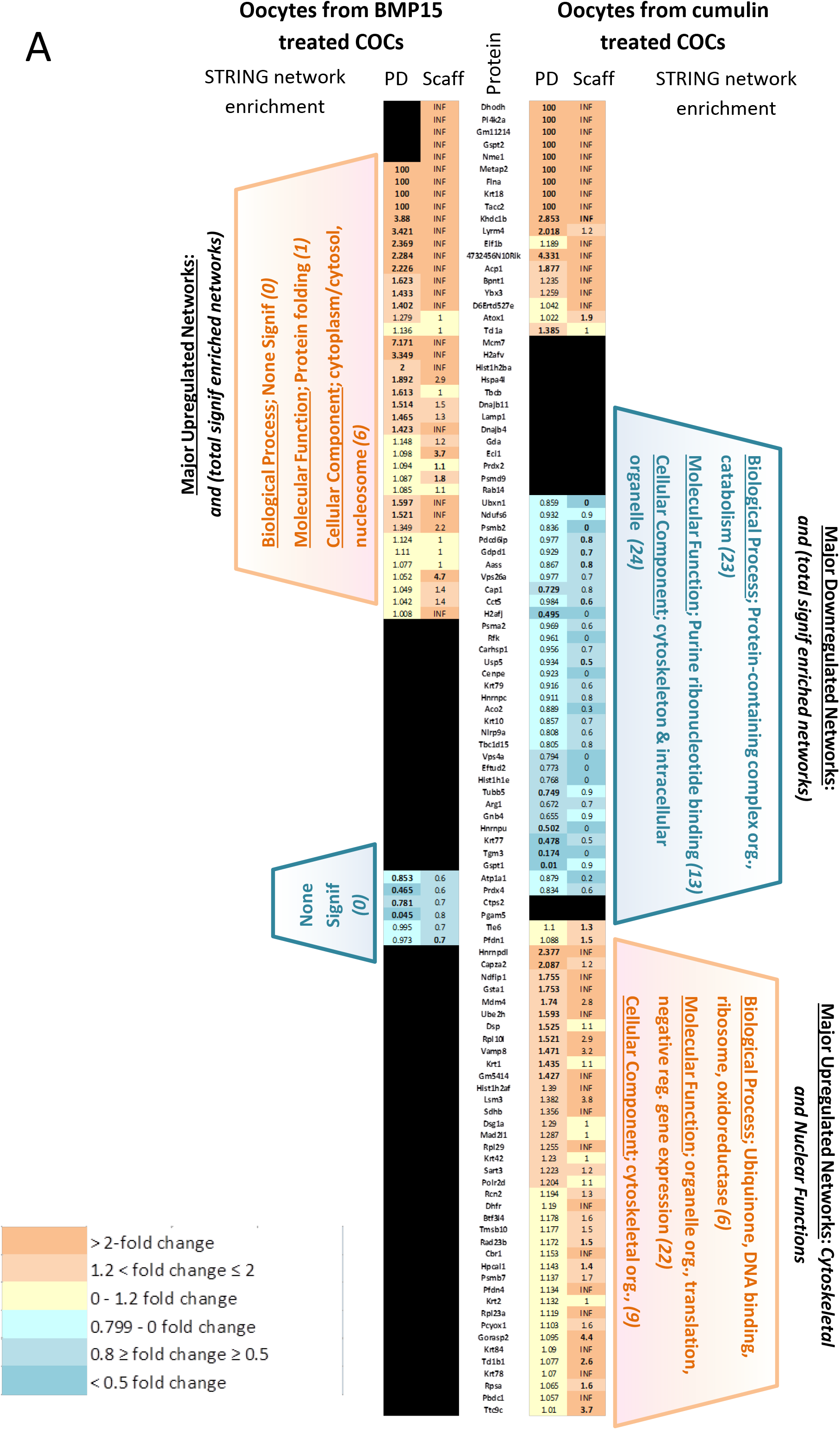
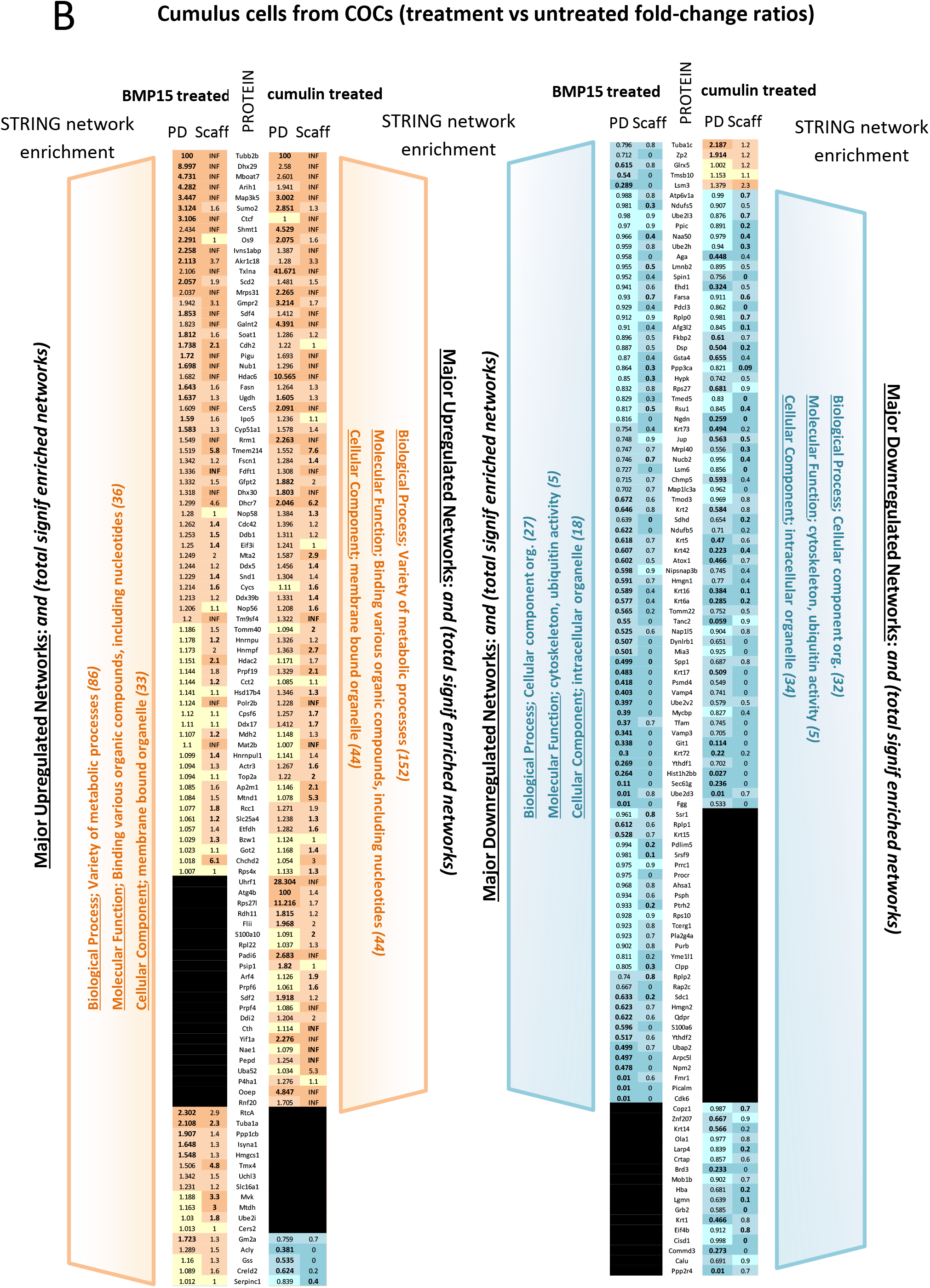
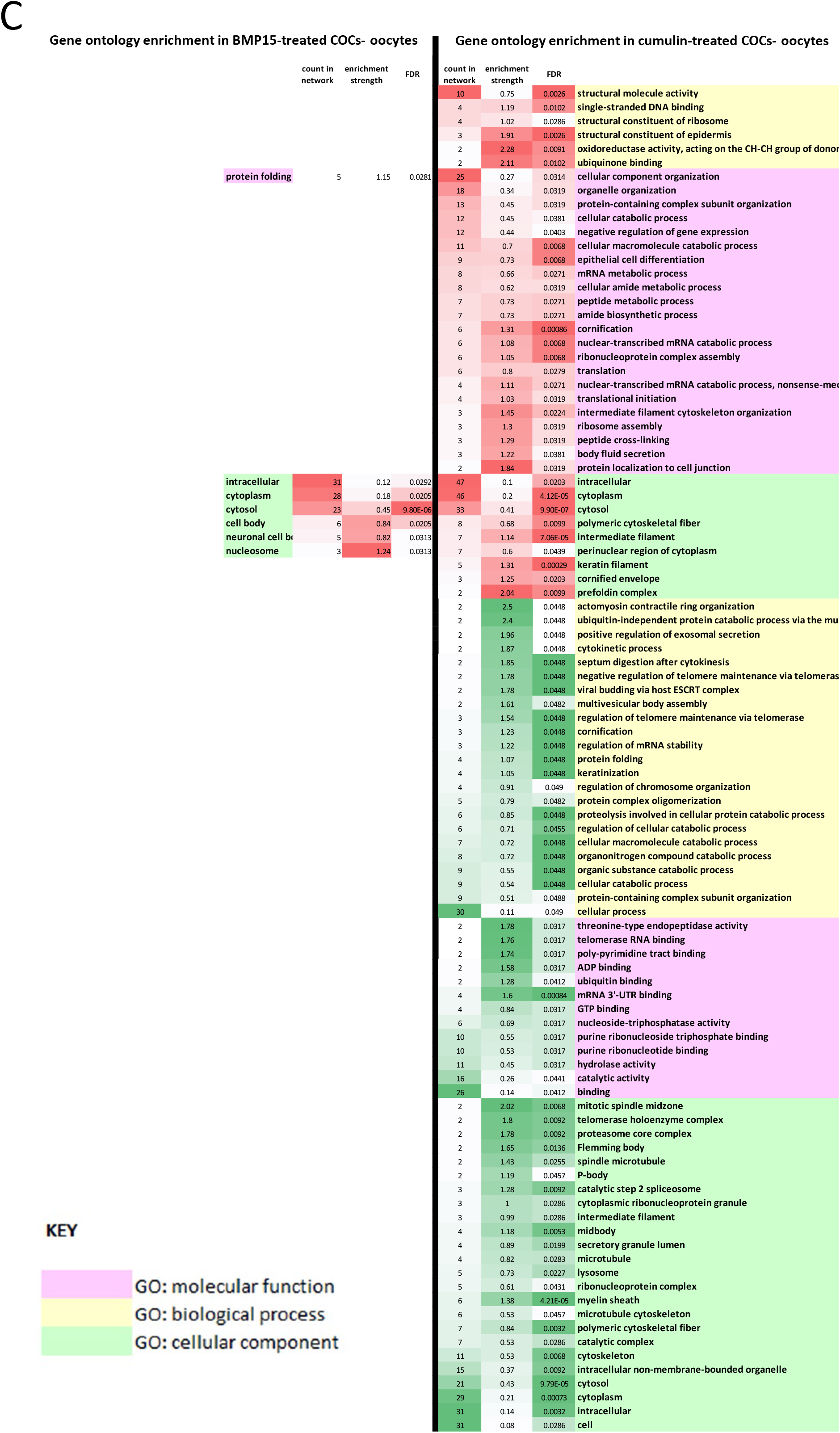
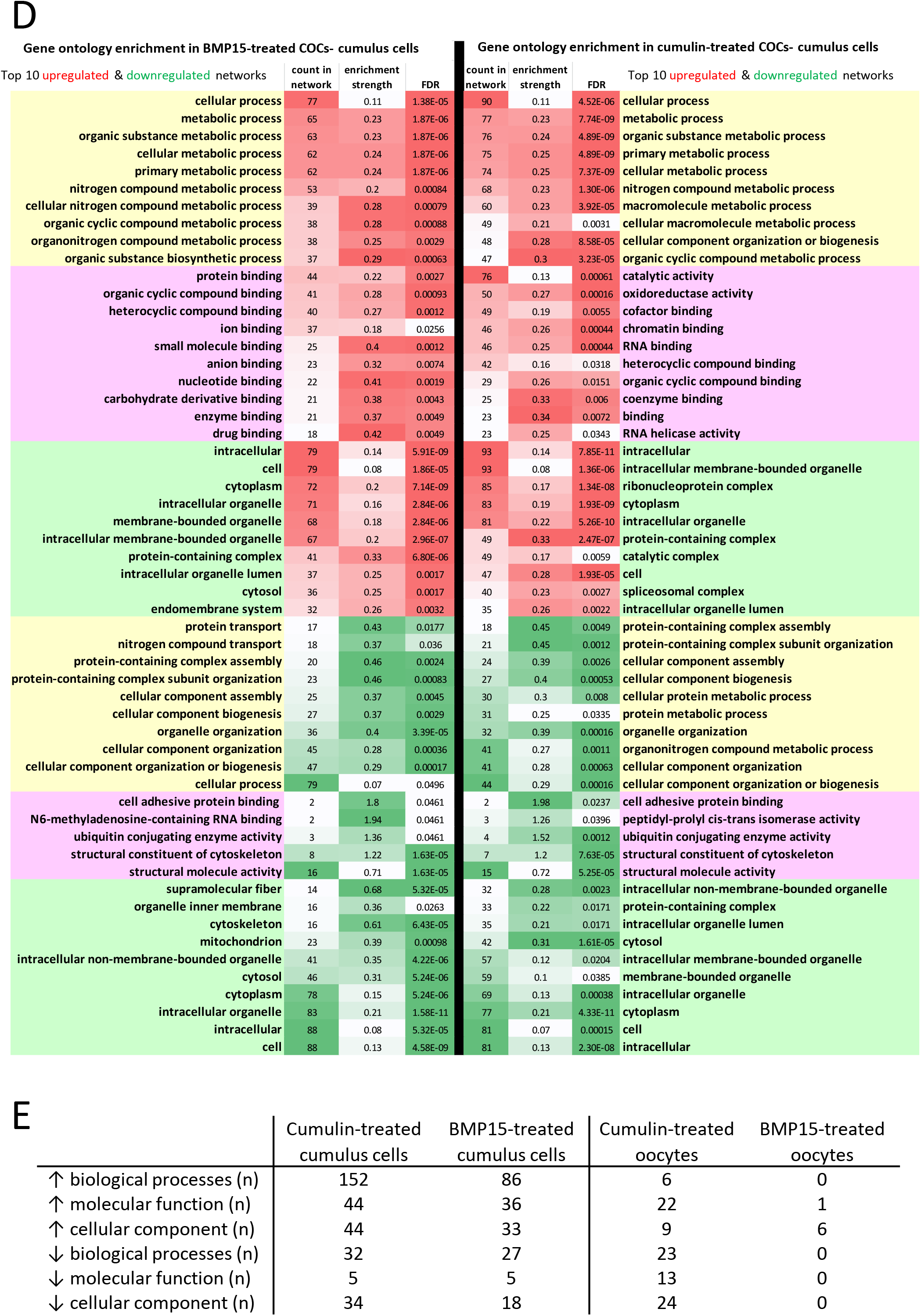
Heat maps of differentially expressed oocyte (A) and cumulus cell (B) proteins identified and quantified in BMP15 and cumulin treated COCs across two different proteomic software analysis platforms; ProteomeDiscoverer v2.4 (PD) and Scaffold v 4.11.0 (Scaff). In the Scaffold data sets, proteins which are not identified in the control samples result in a zero value for the ratio denominator and are marked as “INF”. In PD2.4 proteins which are not identified in the control samples and result in a zero denominator are nominally dubbed with an arbitrary high value (*i*.*e*., 100). All proteins were identified with high confidence across both platforms and included identification with a minimum of two peptides. To combine the two quantification datasets (PD2.4 peak area ratio and scaffold normalized spectral count ratios) for the process of enrichment analysis, those proteins were selected for which the ratio changed in a consistent direction across both platforms, and where at least one platform registered at least a 20% change and/or had a statistically significant difference (p<0.05). Regression analysis data comparing peak area ratio vs normalized spectral counts ratio data is shown in supplementary Figure S1. String network enrichment of biological processes, molecular functions and cellular components for up and down regulated proteins are shown either side of the heat maps, using matching color for up and down regulation (i.e., brown and blue respectively). A heat map representation of enrichment in specific gene ontology (GO) networks following COC treatment with BMP15 (left side) or cumulin (right side) for oocytes (C) and cumulus cells (D). The significantly enriched network lists for oocytes are shown in full, while the top 10 networks for cumulus cells are shown, which subsume the highest number of proteins are displayed in (D). Total significantly enriched network numbers across treatments and across cell types are shown in (E).

### Effects of BMP15 and cumulin treatment of COCs on cumulus proteome expression

The effect of BMP15 or cumulin treatments of COCs resulted in much greater proteomic change in cumulus cells than in oocytes, even taking into consideration the different protein loadings originally used. The different protein loadings resulted in identification of about double the number of proteins in cumulus cells than in oocytes (Fig. 3A). However, there were 4-5 times as many differentially expressed proteins in cumulus cells than in oocytes (Fig. 3A).

Cumulin appeared to have a greater impact on proteomic expression in both cell types than BMP15 did, however this difference was quite marked in oocytes, and less so in cumulus cells. There was also limited overlap in the specific proteins that were differentially expressed in response to each treatment, in oocytes (Fig. 4A), while there was considerably greater overlap of specific up- and down-regulated proteins in cumulus cells in response to both BMP15 and cumulin (Fig. 4B).

Network analysis showed that cumulus cells had a mixture of biological and functional networks, which were affected by BMP15 and cumulin exposure of COCs (Figs. 4B, 4D, 4E); for cumulin treated cumulus cells, there were 240 and 60 networks in the up- and down-regulated protein lists, respectively, with roughly a third fewer networks in the BMP15 treated list (Fig. 4E). Interestingly, when the top 10 networks were compared in heatmap format (Fig. 4D), there is a remarkable similarity in cumulus cell responses to BMP15 and cumulin. Notably, the top ten upregulated biological processes represented in the network analysis were almost exclusively diverse metabolic processes, though mostly non-mitochondrial, and instead based in the cytoplasm (Fig. 4D). Other upregulated molecular functions include oxidoreductase activity and chromatin and RNA binding. As with the BMP15 treatment, the cumulin downregulated cumulus cell proteome largely revolved around cytoskeletal and organelle organization.

### Effects of BMP15 and cumulin on organelle ultrastructure and content

Transmission electron microscopy revealed that cumulin altered cumulus cell mitochondrion and endoplasmic reticulum morphology (Fig. 5A). Relative to untreated controls, cumulus cells of cumulin-treated COCs exhibited more rounded and swollen mitochondria, and more dilated endoplasmic reticulum (ER) with wider cisternae, implying a higher workload for both organelles. BMP15-treated cumulus cell mitochondrial morphology more closely resembled control cells, with regular bean-shaped mitochondria with regular cristae, however slight dilation of the ER cisternae was evident. In oocytes, the control group exhibited the oocyte-typical round, hooded mitochondria with thinner outer membranes, while cumulin-treated oocyte mitochondria exhibited a more swollen outer membrane (Fig. 5A).

**Figure 5:**
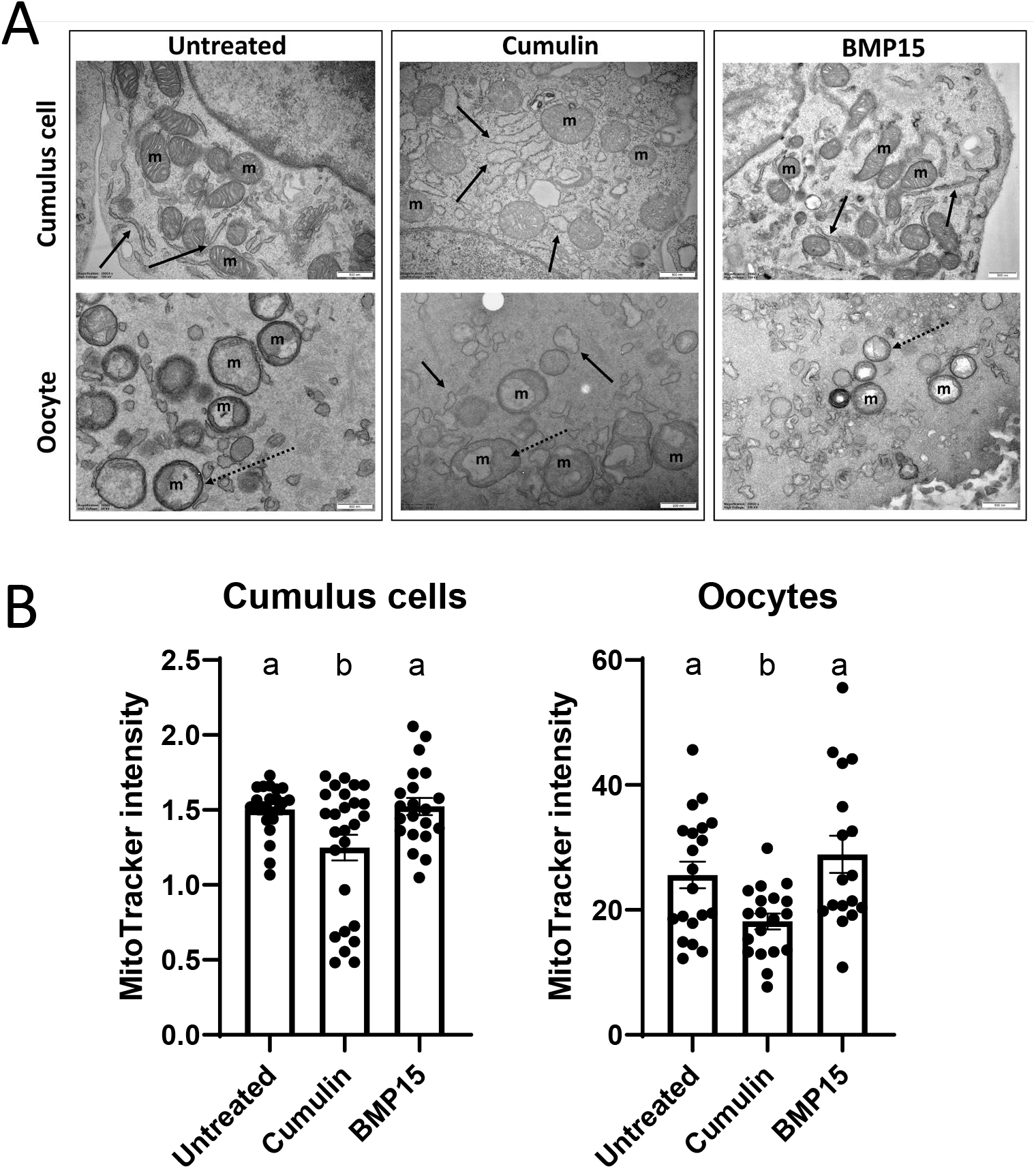
(A) Transmission electron micrographs of oocytes and cumulus cells from BMP15 and cumulin treated COCs, with solid arrows highlighting endoplasmic reticulum in each treatment condition, example mitochondria labelled “m”, and dashed arrows indicating swollen outer membrane of mitochondria. (B) MitoTracker staining used to estimate the numbers of mitochondria in oocytes and cumulus cells from BMP15 and cumulin treated COCs. Bars represent means ± SEM, N=17-20 oocytes collected over 3 biological replicate experiments. For cumulus cells, averaged readings 4-10 cells per COC over 21-26 COCs, collected over 3 biological replicate experiments. Bars with no common superscripts are significantly different (p<0.05; one-way ANOVA with Tukey’s post-hoc test).

To explore whether mitochondrial numbers were also affected, a MitoTracker staining technique was used as a proxy for mitochondrial counts (Fig. 5B). A significant decrease of ∼20-25% in pixel intensity was observed in cumulin treated COCs (in both cell types), relative to control (Fig. 5B), which is likely indicative of lower overall mitochondrial numbers and/or functionality (membrane potential) in cumulin treated cells. There was no effect of BMP15 in either cell type.

### Effects of BMP15 and cumulin on cellular respiration

Given the notable absence of mitochondrial protein enrichment in the upregulated protein network analysis, despite extensive enrichment of a diversity of metabolic pathways, together with alteration of several mitochondrial proteins (Figs. 4A, 4B, 6A), respiration was investigated in COCs following IVM ± cumulin or BMP15. Relative to untreated controls, the basal oxygen consumption rate (OCR) was significantly lower in cumulin-treated COCs, but not BMP15 treated COCs (Fig. 6B). To measure maximal mitochondrial respiratory capacity, COCs were exposed to mitochondrial inhibitors following basal OCR measurement, allowing the measurement of maximal OCR. Relative to untreated controls, maximal OCR was significantly lower in COCs exposed to cumulin, but not BMP15, during oocyte maturation (Fig. 6C). These data support the observation of reduced pixel intensity in the MitoTracker assay (Fig. 5B) and are consistent with either reduced mitochondrial numbers or functionality following cumulin treatment.

**Figure 6:**
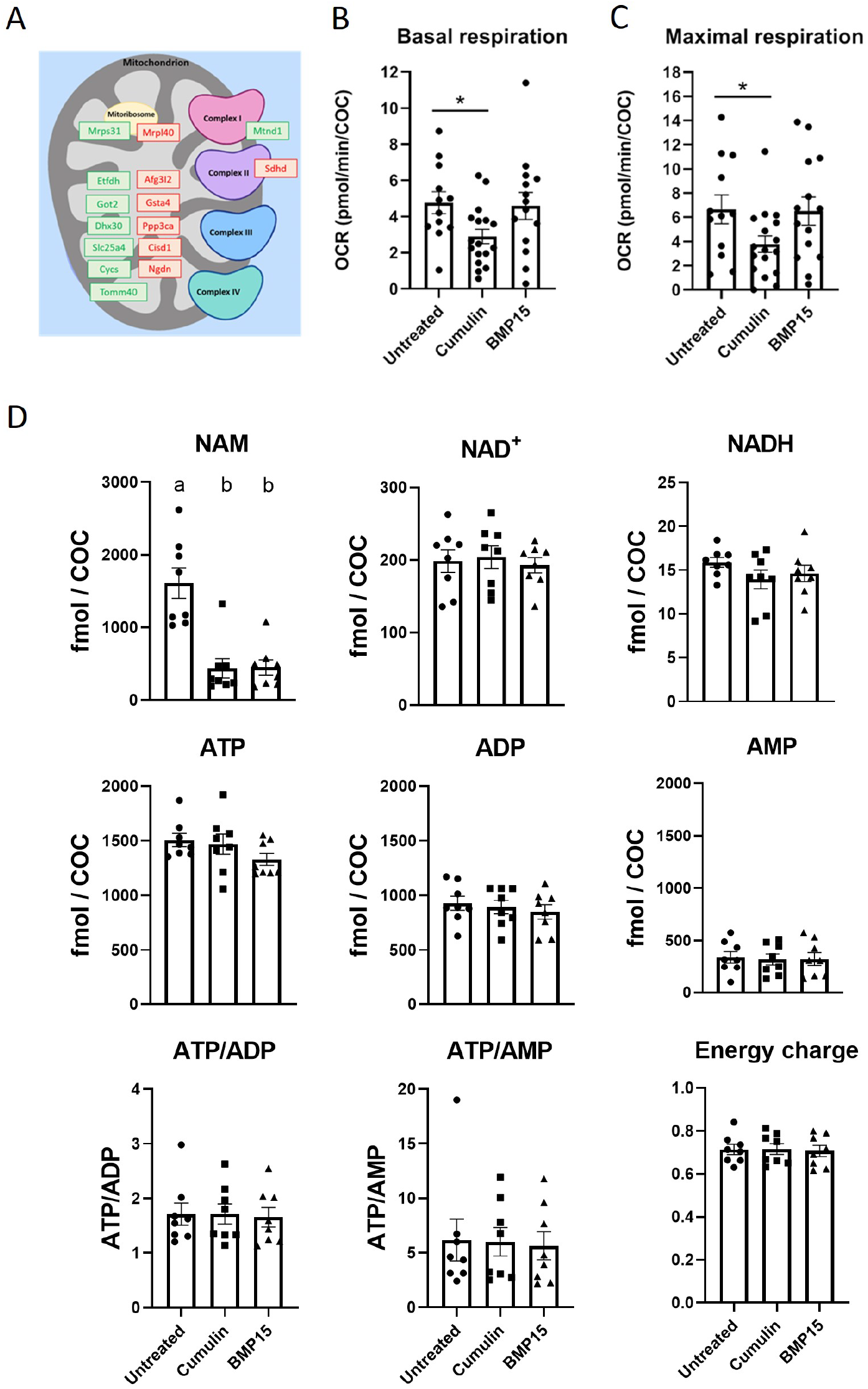
Functional studies of cellular metabolism. (A) Mitochondrial proteins differentially expressed in cumulin-treated (but not BMP15-treated) cumulus cells suggested that cumulin may alter mitochondrial function (green = upregulated proteins, red = downregulated proteins). COC basal (B) oxygen consumption rate (OCR) and maximal OCR (C) were measured to assess mitochondrial activity. Bars present mean ± SEM (representative of 12-17 wells containing 5-15 COCs each, measured over 4 replicate experiments. Each datapoint on graph represents a measurement from one well. Data is normalized per COC in each well. * p<0.05 Mann-Whitney test between the two groups. (D) NAD^+^ and adenosine metabolites were measured in COCs. Effect of cumulin and BMP15 on NAD^+^ and adenosine metabolites in COCs. Metabolite levels were quantified in BMP15 and cumulin treated COCs. Bars present mean ± SEM. N= 8 biological replicates. Data is normalized per COC. Different superscripts demote a significant difference, one-way ANOVA, p<0.05.

### Effects of BMP15 and cumulin on adenosine and NAD^+^ metabolites

Since both the MitoTracker assay and respiration data suggested decreased mitochondrial function in response to cumulin, the energy state of the COCs was examined by assaying the levels of adenosine and nicotinamide nucleotides (Fig. 6D). COC adenosine nucleotides ATP, ADP, and AMP, the ATP/ADP ratio, and ATP:AMP ratio, were not statistically different across treatments, and cellular energy charge was unaltered by exposure to cumulin and BMP15 (Fig. 6D). From the NAD^+^ metabolome, NAD^+^ and NADH were unaltered by cumulin or BMP15, however nicotinamide (NAM) was markedly and significantly decreased by both cumulin or BMP15 (Fig. 6D). Together, these data show that energy balance is maintained in the COC, despite lower mitochondrial number (mitotracker, Fig. 5B) and respiration (Seahorse assay, Fig. 6B,C), implying that existing mitochondria may be working harder to maintain energy balance, very likely supported by markedly upregulated cytoplasmic metabolism, which is anticipated to be occurring based on the types of upregulated protein networks identified via proteomic analysis (Fig. 4). Additional workload on existing mitochondria in response to cumulin is supported by the mitochondrial swelling observed in the cumulus cells when viewed by TEM (Fig. 5A).

### Effects of BMP15 and cumulin on cellular spectral profiles

Multispectral discrimination modelling utilised cellular image features from the autofluorescent profiles of oocytes and cumulus cells. The classifier applied to these data achieved a high degree of separation of BMP15 or cumulin treated cells versus untreated cells (intersection of union of 5-22%; Fig. 3D). This provides direct evidence that cumulin and BMP15 have a major impact on the molecular composition and behaviour of oocytes and cumulus cells. Accordingly, native fluorophore spectra of NAD(P)H, flavins, and cytochrome C were extracted (Fig. 7) and the relative abundance of these compounds was calculated. NAD(P)H was significantly lower in the cumulus cells of cumulin- and BMP15-treated COCs, compared with untreated COCs (Fig. 7A). The relative abundance of flavins was increased by cumulin but decreased by BMP15 (Fig. 7A). Cytochrome C was significantly elevated by BMP15 but not cumulin (Fig. 7A). The NAD(P)H:flavins ratio, an indicator of cellular redox state, was significantly reduced by cumulin (Fig. 7A). In oocytes, NAD(P)H was reduced by BMP15 and had a tendency to be lower in response to cumulin (Fig. 7B). The redox ratio also tended to be lower in response to both OSFs, albeit non-significantly (Fig. 7B).

**Figure 7:**
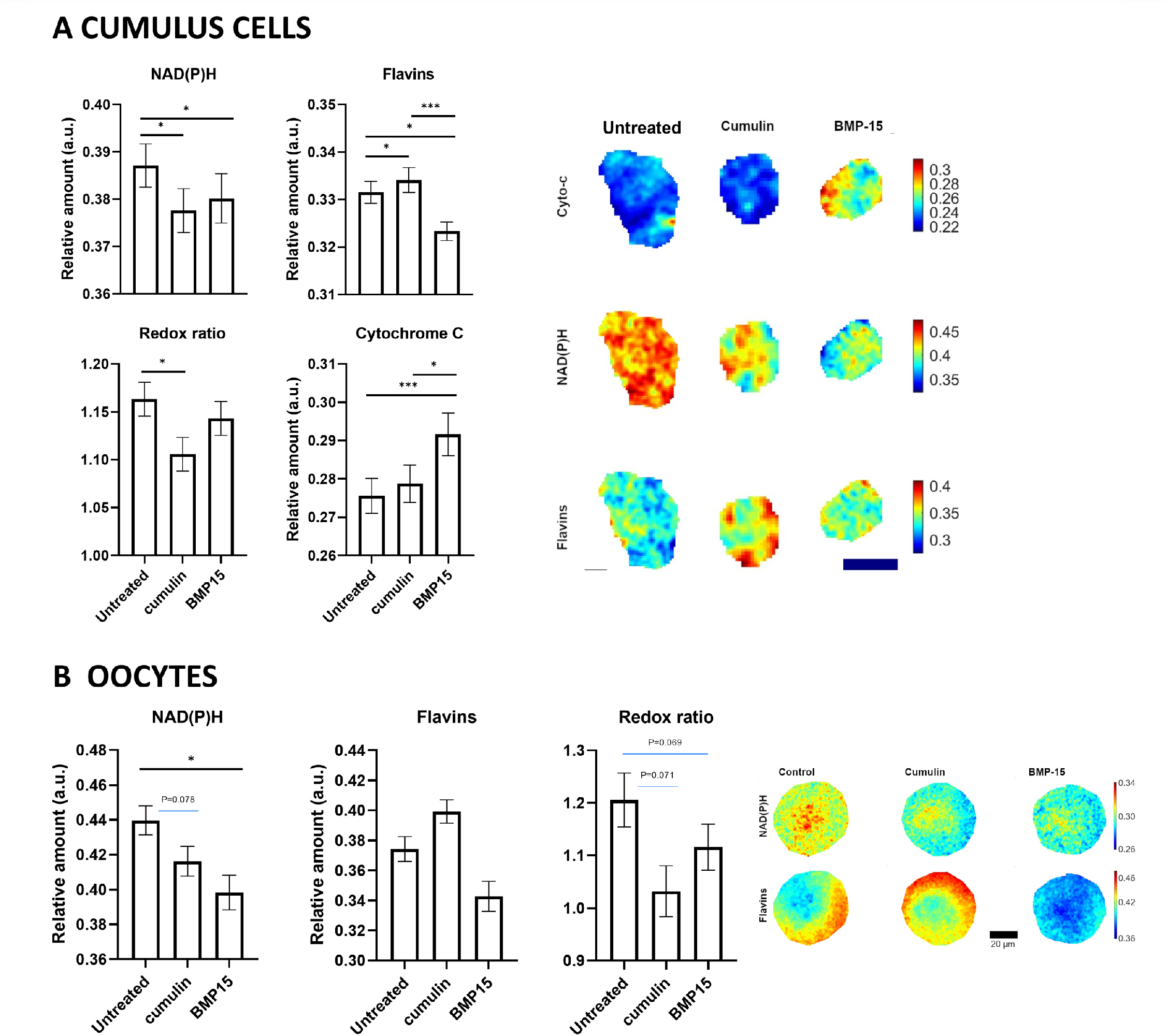
Relative abundance of NAD(P)H, flavins, the redox ratio (NAD(P)H/flavins) and/or cytochrome-C in cumulus cells (A) and oocytes (B) as determined by multispectral unmixing. Images are representative abundance maps of these fluorophores for each treatment group and cell type. Bars represents the median ± SEM of the relative abundance. Normalized abundance scale is used for the visual comparison. Scale bar is 5 µm. N=137-179 COCs analyzed over 4 biological replicate experiments. * (p<0.05), ** (p <0.01) and *** (p <0.001) using two sample t-test.

## DISCUSSION

This study provides a detailed exploration of the mechanisms by which oocyte-secreted factors affect the cooperative (auto-symbiotic) relationship between oocytes and cumulus cells within the COC during oocyte maturation, with consequent improvements of fertility and embryonic development. Two orthogonal global analyses (proteomics and multispectral analysis) showed that the cellular profiles of untreated oocytes and cumulus cells were clearly differentiated from OSF treated cells (Fig. 3). Cellular morphology using TEM revealed morphological changes to organelles in both cumulus cells and oocytes, particularly mitochondria and ER structure. Finally, specific functional studies focused on changes to cellular energetic profiles were performed, since data from the global analyses indicated that metabolic processes represented some of the most significantly enriched pathways. Accordingly, oocyte and cumulus cell REDOX states were altered by OSFs. Cumulin also significantly decreased mitochondrial content and activity, however net ATP and energy homeostasis were unaltered. Collectively, these data demonstrate that oocyte paracrine signaling remodels COC metabolism in preparation for ensuing fertilization and embryonic development.

### Cellular cooperativity proceeds by cellular division of labor in support of enhanced fertility

Following OSF exposure, both cell types exhibit distinct autofluorescent profiles, and a differentially altered expression of a subset of proteins. OSF exposure had a greater effect on the cumulus cell than oocyte proteome, with a greater number of differentially expressed networks and proteins elicited in cumulus cells than oocytes. This is likely because cumulin and BMP15 are secreted from the oocyte and their primary target is the cumulus cell, which expresses their target receptors [14], with their effects on oocytes being a consequence of altered cumulus cell function.

In this study, a substantial number of oocyte upregulated proteins were identified as being involved in DNA binding and nuclear function, suggesting that cumulin affects meiosis in the oocyte and may increase meiotic fidelity. By contrast, cumulus cells were characterized by substantially larger numbers of differentially expressed proteins (both up and downregulated), with the most upregulated biological processes being a diversity of metabolic pathways, including upregulation of cytoplasmic metabolic pathways, mRNA splicing and nuclear protein production, and downregulation of mitochondrial proteins. Hence, OSFs may direct a division of labor between the oocyte and its supporting cumulus cells, as originally proposed by Eppig, Wigglesworth [41], but specifically a division which favors minimization of routine cellular activity in the oocyte (with a focus on nuclear functions and minimization of oxidative stress), while meeting its metabolic and building block needs by massively ramped-up metabolic activity within the cumulus cells.

### Oocyte paracrine factors induce metabolic plasticity in COCs

It is well known that the oocyte’s metabolic capacity is limited. It is unable to metabolize glucose into pyruvate for energy and instead relies on cumulus cells to provide pyruvate [42-44], and it lacks the machinery for amino acid uptake and cholesterol biosynthesis, processes which are instead performed for it by cumulus cells [28, 45, 46]. As such, oocyte paracrine control of cumulus cell metabolism is a critical mechanism to compensate for the oocyte’s metabolic deficiencies [9, 28].

BMP15 and cumulin treatment caused upregulation of a variety of cytosol based metabolic pathways in cumulus cells, which are not related to oxidative phosphorylation or mitochondrial metabolism. The majority of differentially expressed metabolic processes were apparent in cumulus cells, where mitochondrial metabolic processes were downregulated, while numerous alternate metabolic pathways such as lipid, nucleotide and carbohydrate based metabolic processes were upregulated. Hence, under a condition where oocyte quality is enhanced, metabolism is redirected away from routine mitochondrial metabolism and towards expending energy to generate building blocks, presumably to supply to the oocyte, to bolster and sustain its stores for the transcriptionally quiescent phase of fertilization and early embryo development.

Proteomic alterations to mitochondrial proteins are suggestive of altered function and are supported by electron microscopy analysis showing morphological changes to mitochondria and the endoplasmic reticulum, while MitoTracker staining indicates decreased mitochondrial numbers in cumulin treated oocytes and cumulus cells. In support of reduced mitochondrial number, the multispectral unmixed data show lower NAD(P)H levels. Energy metabolite data show that most cellular energy pathway metabolites, particularly ATP, remain similar in untreated and treated COCs, and overall energy metabolites are in balance. The notable exception is nicotinamide which is markedly lower in both BMP15 and cumulin treated cells. As the main precursor to NAD^+^ pathway metabolites (including the redox cofactor NADPH), the lower levels of nicotinamide may reflect its higher consumption to maintain cellular homeostasis driven by OSFs, which may occur if metabolism is redirected from respiration to other metabolic pathways.

Having cumulus cells subsume housekeeping roles on behalf of the oocyte may simultaneously provide for their cellular maintenance needs while allowing the oocyte to minimize stress, such as reactive oxygen species generation, which would adversely affect the oocyte genome and, must therefore be avoided for propagation of the species [47]. Such redirection of metabolism to other pathways, indicative of metabolic plasticity, is reported in cancer cells where alternate pathways supporting the synthesis of lipids, proteins and nucleic acids, are upregulated as required for cell growth and proliferation [48, 49]. Krisher and Prather [50] have also proposed that oocytes may utilize a metabolic strategy like the Warburg Effect in preparation for rapid embryonic growth after fertilization, whereby glucose is used to generate glycolytic intermediates for ribose-5-phosphate and NADPH production via the pentose phosphate pathway, and lactate is generated from pyruvate to maintain NAD+ levels to support elevated glycolysis. Metabolic plasticity as a driver of mammalian embryogenesis has also recently been reported [51]. We suggest that metabolic plasticity may also be facilitated by OSFs during oocyte maturation, where such oocyte paracrine factors redirect metabolism of some glucose in cumulus cells (e.g. to the pentose phosphate pathway), to facilitate the synthesis of these macromolecules to support cumulus cell proliferation and build oocyte reserves, rather than to generate ATP via mitochondrial oxidative phosphorylation. This may ultimately serve to support fertilization, embryonic survival and development. Furthermore, metabolic plasticity may also regulate nuclear signaling and epigenetic mechanisms [49, 52]. Supporting this notion, we note that RNA and DNA processing pathways were significantly enriched in cumulin-treated cumulus cells. Hence, our data support the hypothesis that OSFs promote cumulus cell metabolic plasticity, likely contributing to improved subsequent embryonic survival and development.

Adaptations to changes in energy demand can be achieved by modifying the number and morphology of mitochondria, as well as the abundance of certain electron transport chain constituents [53-56]. Cumulin treated cumulus cells exhibited all three of these modifications. TEM assessment of cumulus cells identified morphological changes to mitochondria, which were more rounded and swollen, with fewer cristae relative to untreated cells. Expression of the electron transport proteins Sdhd and Mtnd1, as well as thirteen other mitochondrial proteins, was also significantly altered (Fig. 6A). Accordingly, mitochondrial number, mitochondrial respiration, and redox potential were significantly decreased in response to cumulin. These data indicate that oocyte paracrine signaling can induce cumulus cell mitochondria to enter a state of respiratory quiescence through remodeling of protein expression and morphology. Sieber, Thomsen [56] also showed that the mitochondria of maturing *Drosophila* and *Xenopus* oocytes undergo quiescence via electron transport chain remodeling and propose that this is an evolutionarily conserved aspect of oocyte development. It remains unclear why mitochondrial quiescence prior to fertilization is beneficial. It may be necessary to limit oxidative damage since mitochondria are a major source of reactive oxygen species (ROS), and mitochondrial ROS are thought to be the main driver of the aging process through dysregulation of cellular homeostasis [57, 58]. The oocyte, spindle and DNA are susceptible to ROS damage. Mitochondria are also known to influence different aspects of cellular function by promoting epigenetic modifications. For example, mitochondrial function influences cellular production of SAM-CH_3_, which is involved in the methylation of nuclear DNA [59, 60]. Hence, the mitochondrial quiescence observed in response to cumulin may serve to alter gene expression. Cumulin treated cumulus cells exhibited enrichment of RNA and DNA processing pathways, supporting this possibility. We propose that cumulus cell mitochondrial quiescence, promoted by the oocyte via OSFs, is likely a means to protect the cells from oxidative damage and maintain stored nutrients, thus allowing the redirection of nutrient metabolism to support synthesis of lipids, proteins, and nucleic acids, as discussed above.

Such metabolic plasticity during the final phase of oocyte maturation may be required for the developmental competence of the oocyte as we have previously shown that exposing COCs to cumulin during maturation significantly improves oocyte developmental competence, as evidenced by an increase in blastocyst yield following fertilization [23, 24]. Since cumulin had a greater effect than BMP15 on COC metabolic processes and mitochondrial function, and cumulin significantly out-performs BMP15 in improving blastocyst yield [23], this supports the hypothesis that metabolic plasticity during maturation is a requirement for oocyte developmental competence.

### Differential response of oocytes and cumulus cells to OSFs

Collectively, the proteomic data demonstrate that cumulin exerts a greater effect than BMP15 on the function of both oocytes and cumulus cells. In cumulus cells, the responses to BMP15 and cumulin is qualitatively similar, in that much the same networks are up- and down-regulated by each treatment. Although the pattern of specific networks affected by these two OSFs are similar, the number of proteins per network is higher in cumulin treated cumulus cells, demonstrating that there is a quantitatively stronger response to cumulin than BMP15. Moreover, the difference between BMP15 and cumulin is particularly pronounced in oocytes, where approximately double the number of proteins is differentially expressed in response to cumulin than to BMP15. Additionally, only 7 networks are significantly enriched in BMP15 treated oocytes, while 97 networks are altered in cumulin treated oocytes. The oocyte’s response to cumulin is thought to be entirely directed via cumulus cells, as its receptors are not known to be expressed on oocytes [18]. Hence, the major remodeling of both the cumulus and oocyte proteomes, in particular to alter metabolism, likely accounts for the positive effects cumulin has on oocyte the oocyte developmental program.

Our previous in vitro work has demonstrated that BMP15 predominately activates Smad1/5/8 signaling, while cumulin, comprised of both BMP15 and GDF9 subunits, potently activates both Smad2/3 and Smad1/5/8 signaling [23, 24]. Hence, the greater effect of cumulin on COC protein expression, as well as its distinct expression profile to both untreated and BMP15 treated cells, may reflect cumulin’s additional capacity to activate Smad2/3, and may be the reason cumulin promotes greater COC function and quality than BMP15. Given the differential effects that cumulin and BMP15 elicited on protein expression, mitochondrial function, and, in previous work, oocyte quality [23], greater consideration of the proper cocktail of OSFs is needed in order to optimize future culture systems to generate the best oocytes for assisted reproduction.

Collectively, this study of the impact of OSFs on oocyte and cumulus cell function during oocyte maturation demonstrates that OSFs promote cell cooperativity between the oocyte and its surrounding support cells, the cumulus cells. OSFs play a significant role in remodeling cumulus cell metabolism during maturation, whilst promoting DNA binding, translation, and ribosome assembly in oocytes. The division of molecular effort likely favors minimization of routine cellular activity in the oocytes, with a focus on nuclear functions and minimization of oxidative stress, while the oocyte’s metabolic and building block needs are met by remodeling of metabolic activity within the oocyte’s support somatic cells.

### Grant Support

The study was funded by a grant (APP1121504) awarded to RBG and CAH and fellowships (APP1023210, APP1117538) awarded to RBG from the National Health and Medical Research Council of Australia.

## Conflict of Interests

RBG is a consultant to City Fertility CHA Global on in vitro maturation (IVM) technologies and is a Scientific Advisory Board member to CooperSurgical who sell IVM products. WAS and CAH hold a patent on modifications to cumulin and its application in reproductive medicine.

